# The evolutionary origin of the universal distribution of mutation fitness effect

**DOI:** 10.1101/867390

**Authors:** Ayuna Barlukova, Igor M. Rouzine

## Abstract

An intriguing fact long defying explanation is the observation of a universal exponential distribution of beneficial mutations in fitness effect for different microorganisms. Here we use a general and straightforward analytic model to demonstrate that, regardless of the inherent distribution of mutation fitness effect across genomic sites, an observed exponential distribution of fitness effects emerges naturally in the long term, as a consequence of the evolutionary process. This result follows from the exponential statistics of the frequency of the less-fit alleles *f* predicted to evolve, in the long term, for both polymorphic and monomorphic sites. The exponential distribution disappears when the system arrives at the steady state, when it is replaced with the classical mutation-selection result, *f* = *μ*/*s*. Based on these findings, we develop a technique to measure selection coefficients for specific genomic sites from two single-time sequence sets. Our results demonstrate the striking difference between the distribution of fitness effects observed experimentally, for naturally occurring mutations, and the “inherent” distribution obtained in a directed-mutagenesis experiment, which can have any shape depending on organism. Based on these results, we develop a new method to measure fitness effects of mutations for each variable residue based on DNA sequences isolated from an adapting population at two time points. This new method is not sensitive to linkage effects and does not require one-site model assumptions.

## Introduction

Evolutionary dynamics of a population of nucleic acid sequences is controlled by several acting forces, including random mutation, natural selection, genetic drift, and linkage decreased by recombination. Of central interest is the adaptation of an organism to a new environment, which occurs due to fixation in a population of rare mutations that confer a benefit to the fitness of the organism (Imhof and Schlotterer 2001; Kassen and Bataillon 2006; Acevedo, et al. 2014; Stern, et al. 2014; Wrenbeck, et al. 2017). The existing models with directional selection and adaptation in a multi-site population demonstrate that only those beneficial mutations that are established in a population, as opposed to those becoming extinct, contribute to the average speed of adaptation in the long term. The advantage of each favorable mutation is measured by the relative change it causes in genome fitness (average progeny number). Thus, the knowledge of fitness effects for different mutations is essential for predicting the evolutionary trajectory of a population, such as occurs during adaptation (for example, during the development of resistance of a pathogen to treatment or the immune response). Therefore, a great effort has been invested in their estimates.

In HIV genome, the average-over-genome fitness effect of a beneficial mutation *∽*1% was first estimated using genetic samples from infected patients (Rouzine and Coffin 1999). Finding out the Distribution of the Fitness Effect of mutation (DFE) over genomic sites in viruses and bacteria requires specially designed and rather elaborate experiments (Imhof and Schlotterer 2001; Kassen and Bataillon 2006; Acevedo, et al. 2014; Stern, et al. 2014; Wrenbeck, et al. 2017). Recently, selection coefficients across the sites of the hemagglutinin gene of human influenza A/H3N2 were estimated by fitting the deterministic one-locus model and its approximate extension for two-loci (Illingworth and Mustonen 2012). The authors fit the model to time-series data on allele frequencies of hemagglutinin (HA) gene of human influenza A H3N2. Next, (Keightley and Eyre-Walker 2007) proposed a method of DFE estimation in mutation-selection-drift equilibrium based on the assumption that DFE has the shape of the gamma distribution. They estimated parameters of gamma distribution from maximization of the likelihood under the assumption that the derived sites are binomially distributed. They also reviewed different types of experiments to estimate DFE and noted a lack of understanding of DFE dynamics. In particular, during evolution, DFE is likely to change in time, which is not in agreement with the assumption of Gillespie-Orr theory of constant DFE (Gillespie 1982; Orr 2003). Thus, all these attempts suggest the need for a more general approach based on evolutionary dynamics, which would not be restricted to a one-locus model (Rouzine, et al. 2001a).

A major complication in predicting evolutionary trajectory and estimating of mutational effects is that the fates of individual alleles are not independent due to clonal interference and genetic background effects (Fisher 1930; Gerrish and Lenski 1998). Another factor is epistasis (Pedruzzi, et al. 2018; Pedruzzi and Rouzine 2019b). Recent advancements in theoretical population genetics provide accurate and general expressions for the speed of adaptation of an asexual population, its genetic diversity, mutation fixation probability, and phylogenetic properties within the framework of the traveling wave theory (Tsimring, et al. 1996; Rouzine, et al. 2003; Rouzine and Coffin 2005; Desai and Fisher 2007; Rouzine and Coffin 2007; Brunet, et al. 2008; Rouzine, et al. 2008; Neher, et al. 2010; Rouzine and Coffin 2010; Hallatschek 2011; Good, et al. 2012; Walczak, et al. 2012; Neher and Hallatschek 2013). These models show that evolution in a multi-site genome can be described by a narrow distribution of genomes in fitness, which slowly moves towards higher or lower fitness. The speed and direction depends on the interplay between selection, mutation, random drift, and linkage effects and recombination. The wave has been observed experimentally in the yeast system (Nguyen Ba, et al. 2019). In all these models, the distribution of fitness effects among mutation sites (DFE) serves as an important input parameter, in addition to the population size, mutation rate, and recombination rate.

In the present work, we propose a rather general approach based on mechanistic description of the slow adaptation of virus far from equilibrium, before its DFE spectrum is measured Our general method allows to measure selection coefficients for specific sites, without being restricted to one-site model approximation, which usually does not work for rapidly changing pathogens. As we mentioned above, linkage of many evolving loci greatly modifies the dynamics evolution and other properties. Our method does not depend on the presence of these effects, and applies both in and outside of the solitary wave regime..

The key to our method is the intriguing fact, that DFE for beneficial mutations has frequently an exponential form. Previous studies in *E*. *coli, Pseudomonas aeruginosa, Pseudomonas fluorescence*, poliovirus show that the rate of beneficial mutation often decreases with their fitness effect exponentially (Fig. 1) (Imhof and Schlotterer 2001; Kassen and Bataillon 2006; Acevedo, et al. 2014; Stern, et al. 2014; Wrenbeck, et al. 2017).

**Fig. 1:**
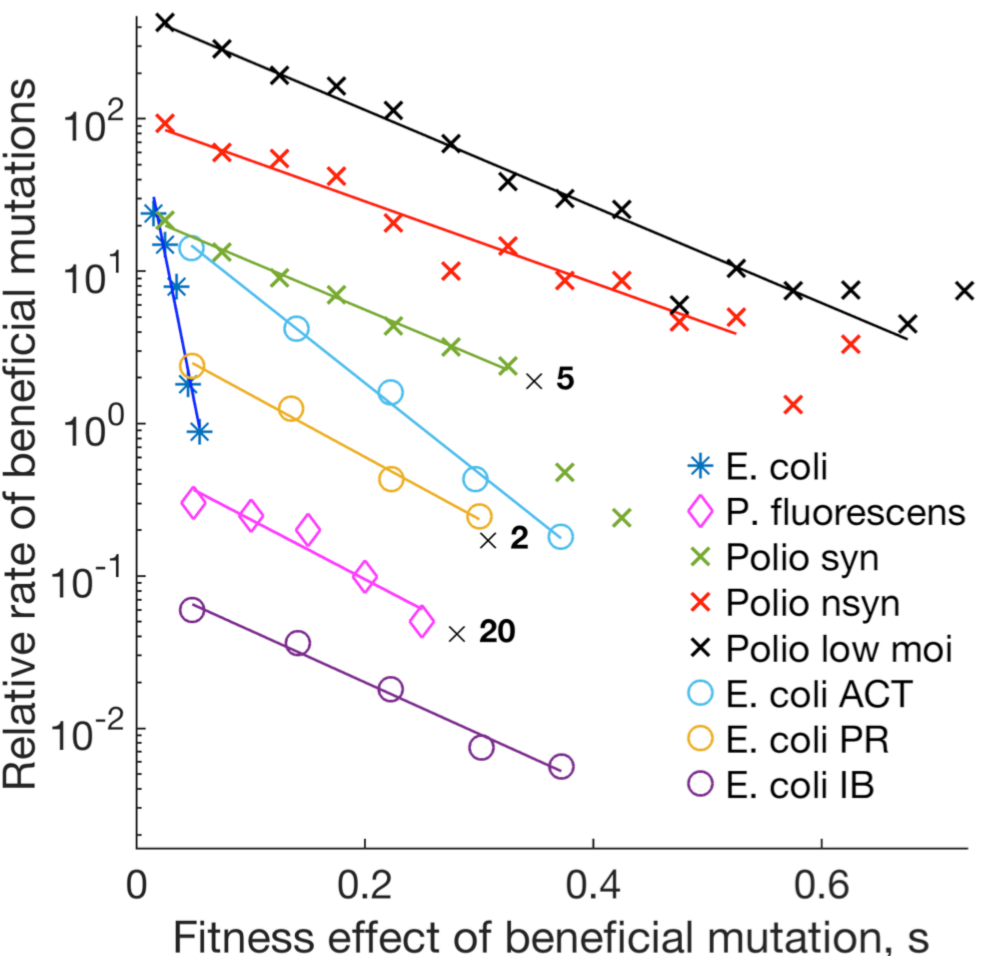
Different studies on distribution of fitness effects of beneficial mutations demonstrate an exponential form. Y-axis: Frequency of beneficial alleles (arbitrary units), DFE(s, t)π(s) in Eq. 1. X-axis: Mutation gain in fitness due to a beneficial mutation (selection coefficient). Symbols represent results obtained for different sites of the genome in experiments on *Escherichia coli* (Imhof and Schlotterer 2001), *Pseudomonas fluorescens* (Kassen and Bataillon 2006), poliovirus synonymous mutations, poliovirus non-synonymous mutations (Acevedo, et al. 2014), poliovirus low MOI (Stern, et al. 2014), *E*. *coli* acetamide (ACT), propionamide (PR), and isobutyramide (IB) (Wrenbeck, et al. 2017).

In the present work, we offer a simple interpretation of this phenomenon. We demonstrate that, regardless of the initial distribution of fitness effects across genomic sites, an exponential DFE emerges naturally, as a consequence of an evolutionary process of slow adaptation. However, the prediction is not completely universal: when the system arrives at a state close to equilibrium, this result ceases to work. Based on these findings to develop a method of estimating the fitness effect of mutation for each variable site in the genome.

We note that, in the existing literature on DFE, two different distributions are referred as DFE. The first is the inherent distribution of selection coefficients of a genome, which represents the genome site density with respect to the values of their selection coefficient. This distribution can be measured directly only by a site-directed mutagenesis experiment and measuring the fitness difference between the two variants for each site (Lee, et al. 2018; Hom, et al. 2019). To avoid confusion, will refer to it as “intrinsic DFE” to emphasize the fact that it is the property of the pathogen/environment and does not depend on the state of population. Another distribution is the distribution of new beneficial mutations arising naturally in an evolution experiment, which depends on the state of adapting population (Fig. 1). We will use term “DFE” to denote the second distribution. We demonstrate below, by mathematical analysis of a general population model in three various regimes, that these two distributions are different from each other. We focus on beneficial mutations only.

## Results

In order to explain the exponential shape of DFE observed in the experiments, we start by noting that beneficial mutations can emerge only at the sites currently occupied with less-fit alleles. Here we assume bi-allelic approximation, when two alleles are considered: the best-fit and the next less-fit. Although each position, in principle, can have four nucleotides A, C, T, G, in real viral data, on moderate time scales 1-10,000 generations, most variable sites display only two alleles in a sample. In this case, if a genomic site is occupied by the less-fit allele, it can become only the best-fit by mutation, and, vice versa, a genomic site occupied by the best-fit allele, it can only lose in fitness. If a population is well adapted during the process of evolution, most of genome sites, in each genome, already carry best-fit alleles and cannot experience beneficial mutations. Therefore, the observed DFE will be affected by the occupation number distribution of less-fit alleles among sites with different *s*, i.e., by the state of population.

Let denote the average frequency of less-fit alleles at a site with fitness effect *s* by *f* (*s*). We note that *f* (*s*) can also be viewed as the frequency of sites available for beneficial mutations. For example, consider a sequence of the form 1000001, where 1 stands for the less-fit allele and 0 for the best-fit allele. Then, only the first and the last positions in the sequence are the sites, where a beneficial mutation can occur, 1 → 0. Thus, the rate of beneficial mutation at any fixed position of the genome must be proportional to the frequency of less-fit allele *f* at this position. If the system is fully adapted, we have *f* = 0, and no beneficial mutations are possible.

### Experiment description

The experiments, shown in Fig 1, consider naturally occurring evolution and count beneficial and deleterious mutations emerging in an adapting population. The authors evolve a population of bacteria or virus for a short time in culture. Newly emerging beneficial mutations result in spontaneous increase in the best-fit allele frequency in time (selection sweeps). Although exact protocols differ, the count occurs for naturally occurring mutations, not for random mutagenesis. In one experiment (Acevedo, et al. 2014), an experimentalist uses a deep sequencing technique CirSeq to monitor the arising frequency of minority alleles at each genomic site as a function of time and fits it with a simple one-site evolution model expression to estimate *s* for each site.,. (Imhof and Schlotterer 2001) focused on beneficial mutations in *E*. *Coli*. They measured selection coefficient *s* for each selection sweep from time series, and then counted the number of sweeps at sites belonging to an interval of the selection coefficient (*X*-axis in Fig. 1). Therefore, all these experiments measure the naturally occurring mutation density, and not intrinsic DFE.

In the last experiment, a beneficial mutation event occurs spontaneously, with a small probability, at a rare less-fit site. If it survives random drift, it gets fixed in the population. We can present the results of these experiments on beneficial mutations (Y-axis, Fig 1) as the product *DFE s, t π*(*s*), where the observed DFE is given by

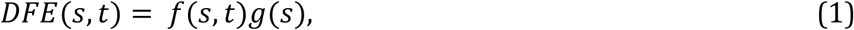

*f* (*s, t*)is the average frequency of target sites available for beneficial mutations averaged over realizations (experimental replicas), *π s* denotes the establishment probability of beneficial mutation *s*, and *g(s)ds* is the number of sites with the selection coefficient in interval [*s, s*+*ds*]. We emphasize that all experiments on natural evolution in Fig. 1 detect the established mutations with density *DFE* (*s, t*)*π* (*s*)= *f* (*s, t*)*g* (*s*)*π* (*s*), and not intrinsic density *g s*. Therefore, the raw distribution of selection coefficient across different sites, intrinsic *g*(*s*), is not the same as the observable distribution in Fig. 1, *DFE s, t π*(*s*). The sites which are already best-fit, cannot have beneficial mutation events. *DFE*(*s, t*) evolves in time, because *f* (*s, t*) changes in time, which explains the aforementioned observation by (Eyre-Walker and Keightley 2007). *DFE*(*s, t*) given by Eq. 1 serves as the input density parameter for the models of evolution in the state of adaptation (Good, et al. 2012).

The intrinsic distribution *g*(*s*), is a biochemical property of the virus and the cell culture conditions. However, *g*(*s*) does not depend on time, unlike *f*(*s, t*), nor it depends on whether a site is occupied by a better-fit or less-fit allele. It is expected to vary broadly between viruses and proteins. For example, some proteins may be more conserved and some less, and *g*(*s*) would be shifted towards larger and smaller s, respectively. Parameter *g*(*s*) is measured in experiment by performing site mutagenesis for each site, one by one, and evaluating fitness differences between the wild type and mutant strains (for example, by growth competition experiment) (Lee, et al. 2018; Hom, et al. 2019). Again, *g*(*s*) does not depend on the state of population, only on a virus and the conditions of its propagation.

In contrast, the value in Fig. 1 is measured by actually evolving the virus in a culture and counting naturally arising mutations in an interval of *s*. The two distributions differ, because in the evolution experiment, a beneficial mutation cannot arise if the site is already occupied with a best-fit allele. In other words, DFE depends on the state of population, occupation number *f*(*s, t*), and *g(s)* does not.

Below mutant frequency *f*(*s, t*) is assumed to have pre-evolved before the experiment for a long time, reflecting pre-history of the population under similar conditions, but is not in mutation-selection drift equilibrium yet, i.e., it is not best adapted yet to the conditions of the experiment. The initial genome in each experiment in Fig. 1 is obtained, originally, by sampling from a previous, well-evolved population, close to the best-fit sequence, with deleterious allele occupation number decaying as given by Eq. 6. Then an experimentalist stores the sample in the freezer. Later the virus is thawed and expanded in the culture, and then it evolves and *s* is measured for spontaneous mutations. Hence, the frequency of uniformly deleterious sites in the initial uniform population an experiment genome mimics the occupancy probability in the previous population. We will describe this pre-evolution of *f*(*s, t*) by simulations and analytically. After predicting the form of *f*(*s, t*) we will use it to estimate intrinsic distribution of *g*(*s*) from data.

We will show below that *f*(*s, t*) depends sharply (exponentially) on *s*, and the log slope of the dependence of *f* (*s, t*) on *s* increases linearly in time. The scale of *g*(*s*) in *s* stays constant. Therefore, sooner or later, exponential *f*(*s, t*) will become faster in *s* than *g*(*s*). It is a well-known mathematical fact that an exponential *f*(*s, t*) multiplied by a slower function *g*(*s*) still appears to be an exponential in the log plot. Therefore, the measured DFE appears as an exponential in the log scale, which explains the experimental results (Fig. 1).

### Model

We consider an asexual organism, which evolved for some time but is still far from the mutation-selection equilibrium before the experiment. A haploid population has *N* binary sequences, where each genome site (nucleotide position) numbered by *i* =1, 2, …, *L* carries one of two possible genetic variants (alleles), denoted *K*_*i*_ =0 or *K*_*i*_ =1. Each site (nucleotide position) has one of two alleles: the better-fit (for example, A), or the less-fit (for example, G). We note that beneficial mutations are rare, which is why it unlikely that two occur at the same nucleotide. We focus here on the short-term adaptation to a new constant environment, where the bi-allelic model is a fair approximation.

The genome is assumed to be very long, *L* ≫ 1. Time is discrete and measured in units of population generations. The evolution of the population is described by a standard Wright-Fisher model, which includes the factors of random mutation with genomic rate *μL*, constant directional selection, and random genetic drift. Recombination is assumed to be absent. We do not consider the case with balancing selection, diploid immune dominance,, or selection for diversity. Once per generation, each individual genome is replaced by a random number of its progeny which obeys multinomial distribution. The total population stays constant. To include directional natural selection (we do not consider balancing selection or diploid dominance), the average progeny number (Darwinian fitness) of sequence *K*_*i*_ is set to *e*^*W*^. We consider the simplest case when the fitness effects of mutations, *s*_i_, are additive over sites:

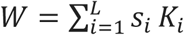

The reference genome, *K*_*i*_ ≡ 0, can be chosen in arbitrary way. For our aim, it is convenient to set it to be the same as the best fit sequence, so that all selection coefficients *s*_i_ are negative. Each site *i* with deleterious allele, *K*_*i*_ = 1, is a target site for a possible beneficial mutation. Vice versa, a site with the favorable allele, *K*_*i*_ = 0, can have a deleterious mutation.

The present model does not consider epistasis and assumes additive contributions of single sites to the fitness landscape. Our rationale for not considering epistasis explicitly is that, in the present work, we consider a short-term dynamics, in which the effects of most epistatic interactions are incorporated in the current values of *s*. If protein evolves for a very long time, a large part of its sequence will change, one mutation triggering changes in other sites, and these changes become permanent and effect the subsequent values of *s* (Rouzine and Coffin 1999; Shah, et al. 2015; McCandlish, et al. 2016). In the long-term, epistasis redefines the values of *s*. Therefore, on very long time scales, when many sites experience replacement of an allele, each site interacts with many other sites. On a short time scale, few sites are polymorphic, and even fewer interact, and not including epistasis implicitly is a fair approximation for most sites. Epistatic interactions with monomorhic sites are embedded in the values of *s*. A more general version of the fitness model that accounts for pairwise epistatic interactions is analyzed in detail in (Pedruzzi, et al. 2018; Pedruzzi and Rouzine 2019c; Pedruzzi and Rouzine 2019a) and, for macroscopic epistasis, in (Good and Desai 2015).

The fitness cost of a deleterious allele *s* is distributed in a complex way among genomic sites. In general, the inherent distribution *g*(*s*) is unknown and depends on a virus, host cell type, and a protein. Its measurement requires an experiment with site-directed mutagenesis along the entire genome. The genome has to be mutated artificially, site by site, and then the value of *s* is measured for each mutation. Below we make no assumptions regarding *g*(*s*) and demonstrate that the exponential shape in the less-fit allele frequency *f*(*s*) arises automatically and independently on the form of *g*(*s*). Later on, we will show how *g(s*) can be calculated from sequence data for the influenza virus and demonstrate that it is an unremarkably slow function within an interval of *s*.

Our work applies only far from mutation selection equilibrium when system is still adapting. It is well known that, in equilibrium, the dependence *f* (*s*) is not exponential, but close to *f* = *μ*/*s* for large *N*. Computer simulations show that the DFE evolves away from an exponential distribution when approaching equilibrium (Eyre-Walker and Keightley 2007).

### Monte-Carlo simulation

We start from an initial population of *N* genomes that has a fraction of deleterious alleles randomly distributed among genomic sites (Fig 2a). Evolution of a sample of hundred sequences in a representative Monte-Carlo run is shown in Fig. 2. For the sake of visual convenience, we have re-ordered genomic sites in the ascending order of the value of selection coefficient *s*_i_.

**Fig. 2:**
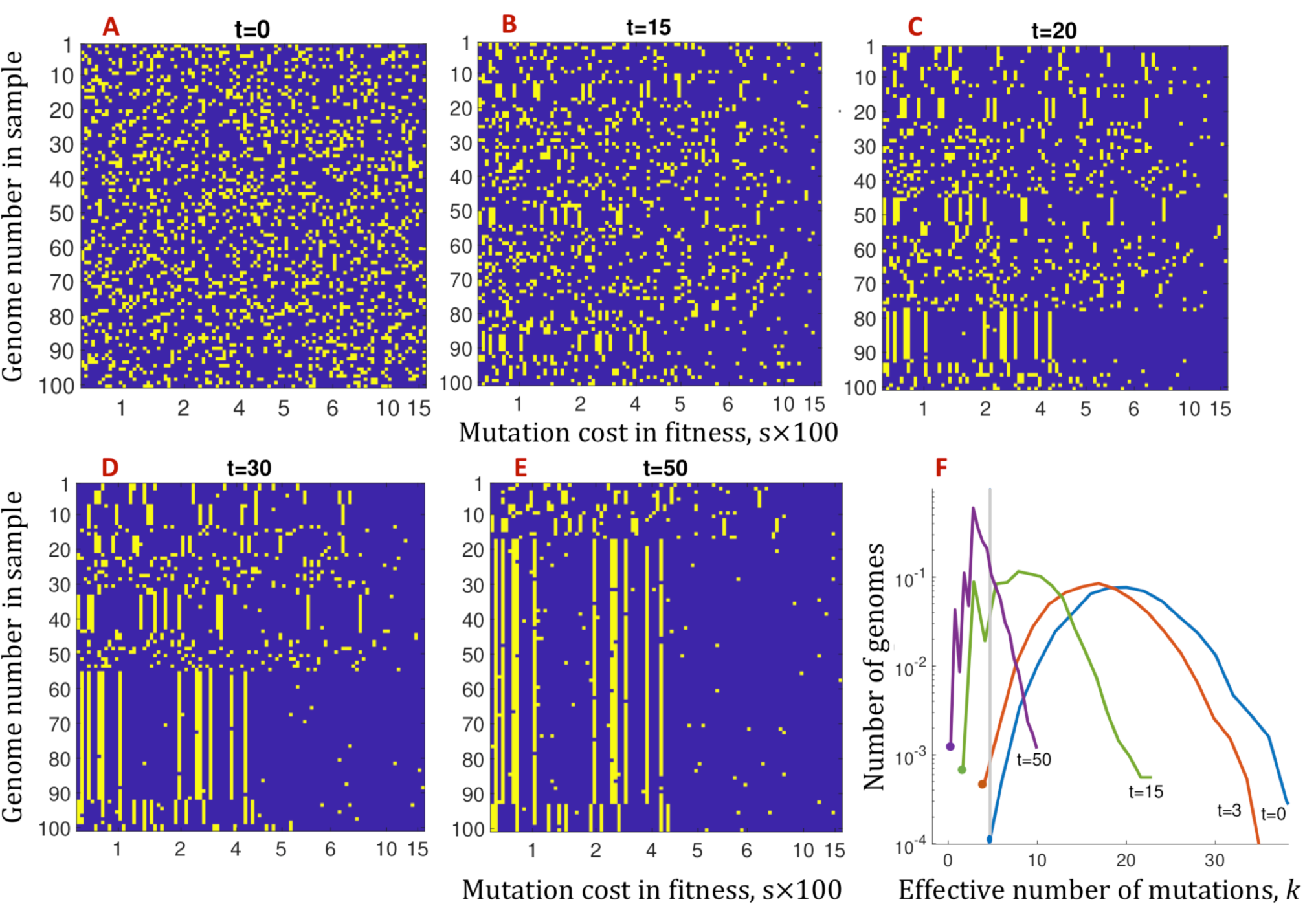
Deleterious alleles with higher values of fitness cost, *s*, are the first to be depleted during the process of adaptation (single run). In other words, beneficial mutations with higher *s* are fixed first. (a-e) Evolution of a sample of 10^2^ sequences. Violet dots: better-fit alleles, yellow dots: less-fit alleles. X-axis: the cost in fitness, *s*, multiplied by 100. The values of *s* are randomly distributed with the half-Gaussian distribution, *s*>0, with the average *s*_*av*_ *=* 0.05. Genomic sites are ordered by the value of *s*. Y-axis: genome number in the sample. The initial population is randomized with the average frequency of deleterious alleles *f*_in_ *=* 0.2. Time points in generations are shown. (f) Evolution of the genome distribution in fitness. X-axis: the effective number of deleterious alleles, defined as *k* = -*W*/*s*_av_. where *W* is fitness. Different colors show discrete time intervals from 0 to 5. Vertical grey line shows the best fitness class of genomes at *t*=0. The emergence of clonal structure in (a-e) coincides with the transition from the selection of pre-existing sequences to the traveling wave regime. Parameters: *f*_in_ *=* 0.2, *N =* 10^4^, *L =* 100, *s*_*av*_ *=* 0.05, *μL =* 0.05. Note that below we study the ensemble average allelic frequency averaged over many runs.

In the process of evolution, we observe increasing redistribution of deleterious alleles among genomic sites as follows (Fig. 2). The sites with a relatively high mutation cost loose deleterious alleles due to natural selection. The asymmetry becomes evident from *t* = 20. Finally, at *t* = 50 (Fig 2e), mutations on the right side are almost absent. Thus, deleterious alleles with higher values of mutation cost vanish earlier, which represents a qualitative explanation of the observed exponential dependence of DFE on *s* (Fig 1).

We note that in our example, we set a rather large value of initial *f*, which is convenient for numerical computations. In real life, mutant frequency *f* may be much smaller than the value we choose. However, our results do not depend on this initial condition assumption. Later, we provide our analytic derivation which is general and applies to very low *f*, as long as they are not in mutation selection equilibrium.

In addition to the observed re-distribution of less fit alleles, we also observe the emergence of group of identical sequences, which we explain by evolutionary process as follows. In Fig 2, two intervals of adaptation can be discerned. Early on, new mutations can be neglected, and the critical evolutionary factor is the natural selection of pre-existing genomes (Fig 2a, b). It was previously revealed by a combination of modeling and experimental evolution of vesicular stomatitis virus (Dutta, et al. 2008). In time interval, *t* ≪ 1/*s*_*av*_, where *s*_*av*_ is the average of *g*(*s)*, the distribution of alleles over genomes remains random.

In contrast, in the second time interval, which starts around *t ∽* 1/*s*_*av*_, where we see formation of deleterious alleles spanning large groups of genomes, new beneficial mutations become crucial for further evolution, because they give birth to new highest-fit genomes (Fig 2b-e). To explain the formation and subsequent growth of groups of identical sequences (Fig 2b-e), we address to traveling wave theory of evolution (Fig 2f).

Formation of these clones occurs at the edge of the traveling wave of fitness distribution (Rouzine, et al. 2003; Desai and Fisher 2007; Hallatschek 2011; Good, et al. 2012) (Fig 2f). The fitness distribution moves in time towards higher values of fitness, i.e., smaller numbers of deleterious alleles. At early times, the distribution is broad and symmetric. In this regime, as was mentioned earlier, the main force is the selection of preexisting genomes. After a while (*t ∽* 1*/s*_*av*_), the profile becomes asymmetric, and the high-fitness edge starts to move to the left together with the peak due to new beneficial mutations (Fig 2f). The genomes, appearing on the left side from the initial high-fitness edge (grey line in Fig 2f) share the initial genetic background. Hence, they produce observed groups of sequences identical at most sites (yellow vertical lines, Fig 2b). As the wave progresses, the clonal structure grows, and eventually, most genomes in the population become an offspring of the same ancestor (Fig 2f).

### Analytic derivation of universal DFE

In this section, we study analytically a general non-equilibrium case of slow adaptation (example in Fig. 2). We assume that the system is very far from steady state, so that *f* is much larger than *f*_*eq*_. We assume also that mutant frequency *f* (*s, t*) has evolved for some time before the experiment measuring DFE, but that the population is not in equilibrium yet, so that deleterious mutation events (reverse mutations) are negligible. Below, we present three independent derivations for three limiting cases, as follows. (i) The case where *f(s,t)* is dominated by polymorphic sites (short term evolution, Fig. 2A,B), (ii) the case where it is dominated by less-fit monomorphic sites (moderate-term evolution, Fig. 2D,E), and (iii) the general case, where both components can be important. In all three derivations, <*f(s,t)* = averaged over realizations is found to be an exponential in *s*, which shrinks in time (see data in Fig. 1). We calculate the exponential slope as a function of time. For details, see Supplement online.

#### (i)Short term evolution

As the above simulation shows, with polymorphic sites at the initial conditions, the evolution of genomes occurring at short times *t* ≪ 1/*s*_*av*_ is mainly due to the selection of preexisting variation and new mutations are not important (*Methods*). Almost all sites are polymorphic due to the polymorphic initial conditions. The probability of having a deleterious allele at a polymorphic site with mutation cost *s* at time *t* has the form

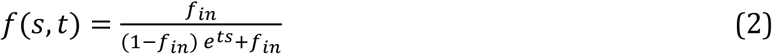

where *f*_in_ is the initial mutant frequency. Note that this expression refers to the ensemble-average frequency. In fact, there are fluctuations between realizations. The slope of the distribution of deleterious alleles is defined as

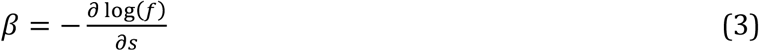

We observe that the formula in Eq 2 does not depend on the initial distribution of selection coefficients among sites, *g*(*s*). At a small initial mutant frequency *f*_*in*_, the formula can be approximated with an exponential, *f*(*s, t*) ≈ exp(-*ts*). The exponential slope is approximately equal to time, *β* = *t* (see Fig. 3). This is an early regime where the evolution of different sites is effectively independent.

**Fig. 3:**
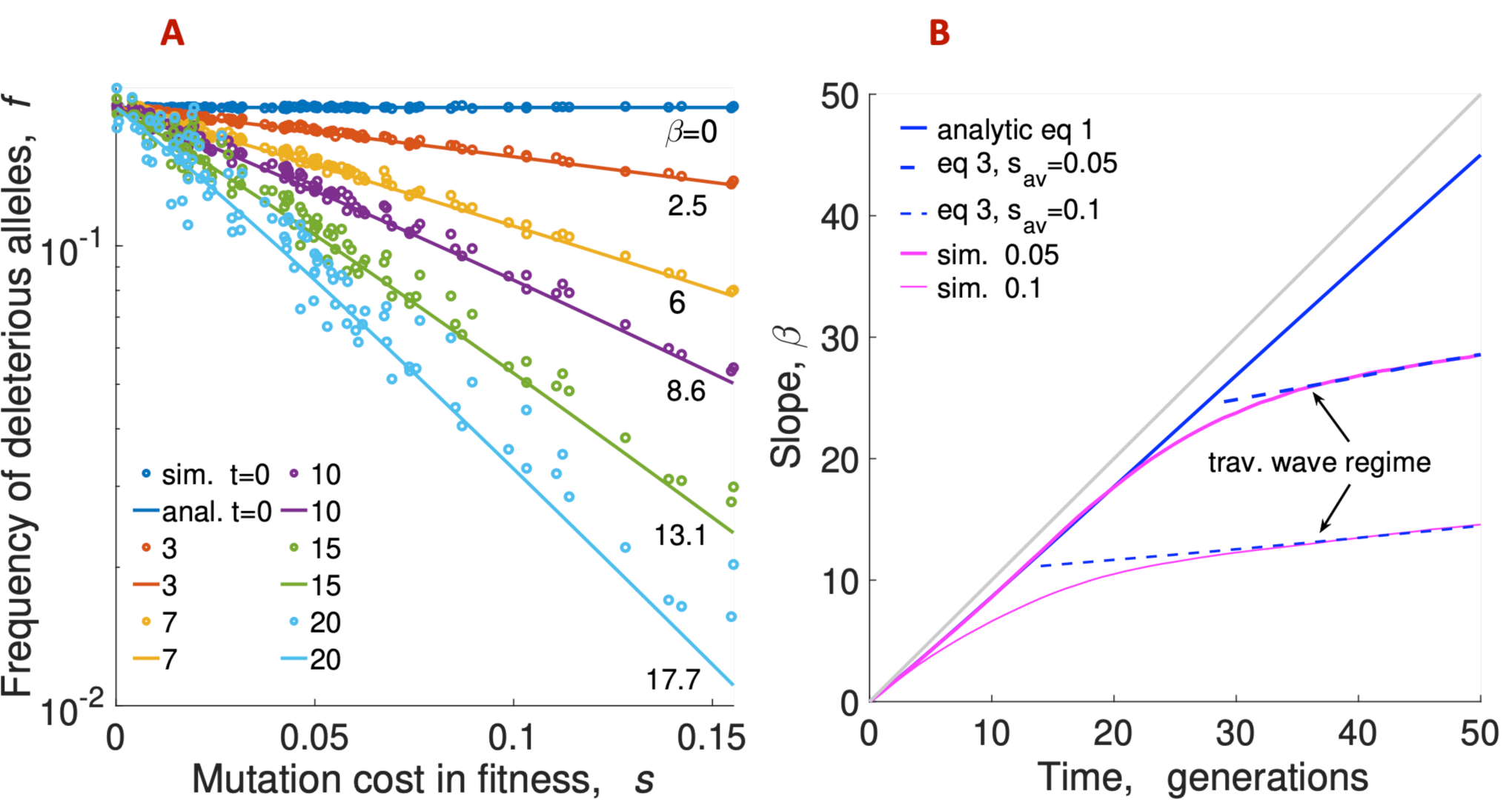
The frequency of deleterious alleles decays exponentially with their fitness effect, with the slope increasing in time. (A) Analytic prediction (Eqs. 1) for the frequency of deleterious alleles agrees with Monte-Carlo simulation. X-axis: Mutation cost of deleterious allele at a genomic site, *s*. Y-axis: Frequency of deleterious alleles at such a site, *f*(*s*). The mutant frequency *f* is averaged over 20 random simulation runs, the straight lines are linear regression. Different colors show different times, symbols are simulation, and lines are analytic prediction (Eq. 1). The numbers on the curves are the values of the slope. Parameters as in Fig 2. (B) The slope of the distribution of deleterious alleles *β*, analytic (blue lines) and simulation (purple lines), as a function of time, *t*. Parameters *f*_in_ and *s*_*av*_ different from those in Fig 2 are shown on the legend. The log-slope for the simulated curves of mutant frequency in (A) is obtained by an exponential fit. We observe that the deviation of the simulated slope from the analytic prediction Eq. 1 at long times coincides with the establishment of the traveling regime, which occurs later for smaller *s*_*av*_ (Fig. 2f). At long times, the traveling wave prediction Eq. 3 applies (dashed blue lines). Grey diagonal shows *β* (*t*) = *t*. Parameters are as in the legend of Fig. 2.

#### (ii)Long term

At longer times *t*>1/*s*_*av*_, beneficial mutations become essential, we jave deleterious alleles for large clones of genomes, and the above derivation does not apply. We need to use the results of the traveling wave theory. In the stationary regime of traveling wave (Fig 2f), fixation of new beneficial alleles is the process that dominates the loss of deleterious alleles (Rouzine, et al. 2003; Desai and Fisher 2007; Hallatschek 2011; Good, et al. 2012). The wave is narrow, most sites are monomorphic, either in less-fit or better-fit allele, and the average allelic frequency<*f*(*s, t*)>is dominated by less-fit monomorphic sites.

Let *t*_0_ be the characteristic time when the traveling wave regime starts. In *Supplementary Methods*, we solve a dynamic equation for the ensemble-averaged allelic frequency, *f*(*s, t*), and obtain

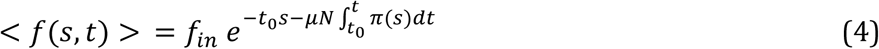

where *π* (*s*) is the probability of fixation of a beneficial mutation with fitness gain *s* derived previously (Good, et al. 2012). The cited paper assumes that an established allele, which does not become extinct, is always fixed. By expanding the argument of the exponential in Eq. 4 in *s*, the slope takes the form

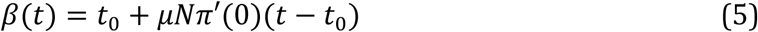

As we can observe, the exponential slope, Eq. 5, depends on time, which is a consequence of the process of slow adaptation [see review (Eyre-Walker and Keightley 2007)]. Please note that Eq. 4 for adaptation regime neglects deleterious mutation events and is valid only far from equilibrium.

#### (iii)General case

Previously, a more general argument was used to predict the exponential shape of the DFE (Pedruzzi, et al. 2018). It does not depend on the knowledge whether deleterious alleles come from monomorphous or polymorphous sites and employs ensemble-averaging for<*f*(*s, t*) >. The main assumption is that, because the wave is slow, the system is in quasi equilibrium, where all the variables change slowly adjusting to the slow change of the average fitness in time. Hence, given the fitness distribution of genomes, the distribution of alleles over sites and genomes is given by the condition that the entropy is in the maximum. However, the full equilibrium does not occur until much later on scaled much larger than 1/<*s*>.

Further, the fitness distribution is narrow, as follows from traveling wave theory, Δ*W* ≪ *W*. Therefore, the system entropy is at the conditional maximum *S* restricted by the average value of fitness -*W*. From these assumptions, we obtained the probability to have a deleterious allele at a given site

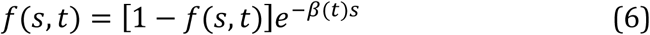

where 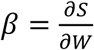. Eq. 6 is general, while Eqs 2 or 5 provide the explicit expressions for β(*t*), which,we remind, depends on time, since the population is not in the true equilibrium but in the process of adaptation.

We note that the mathematical form of Eq. 6 well-known for equilibrium systems. It often appears in biophysics for a two-state equilibrium system connected to a large thermal reservoir (Berg, Lassig, et al. 2004). For the first time, it has been obtained 100 years ago for Fermi statistics. More recently, it was re-derived for the steady state for similar model in steady state *f* = *f*_*eq*_, with *β* = *2N* where *N* is the effective population size (Rice, et al. 2015).

Here we show, for the first time, that the same universal expression can apply far from equilibrium *f* ≫ *f*_*eq*_, when deleterious mutation effects can be neglected, and the system is in adiabatically slow evolution. The expression for β(*t*) given by Eq. 5 is quite different from the equilibrium case (Rice, et al. 2015) and depends on time, which is confirmed by simulation (Fig. 3A). This time dependence, as we said has been observed experimentally (Eyre-Walker and Keightley 2007).

The reason for the non-trivial validity of Fermi statistics far from equilibrium is the adiabatic (quasi-equilibrium) regime which exist in a broad parameter region in the traveling wave regime (Pedruzzi, et al. 2018; Pedruzzi and Rouzine 2019a; Pedruzzi and Rouzine 2019c) This regime applies even for slowly changing selective pressure, such as exists in the case of influenza due to accumulation immune memory cells. This case can be mapped onto traveling wave for constant selection condition (Rouzine and Rozhnova 2018; Yan, et al. 2019). We also point out that, in equilibrium, Eq. 6 does not apply in in the steady state in the deterministic limit of very large *N*. In this case, the correct expression in this case is the famous single-locus formula *f*(*s*) = *μ*/*s*, where *μ* is the deleterious mutation rate [for review, see (Rouzine, et al. 2001b)]. The reason for the failure of Fermi statistics in the deterministic steady state is that deleterious mutations mix the two states [which fact is neglected in the cited work (Rice, et al. 2015)]. Deleterious mutations can be neglected far from equilibrium, *f* ≫ *f*_*eq*_.

Thus, Eqs. 2 to 4 demonstrate that the exponential dependence *f* (*s*) arises in the course of evolution at any initial conditions after the evolution time *t ∽* 1/*s*_*av*_, and that the resulting exponential slope is robust to the initial conditions. The pre-factor at the exponential depends on the time of pre-evolution and *f*_in_. We assume that the system evolved for time longer than the inverse average *t ∽* 1/ *s*_*av*_ but is not in equilibrium yet.

### Monte-Carlo simulation confirms theory

To test our analytic theory, we compare the frequency of deleterious alleles obtained by analytic prediction (Eq. 2) *f*(*s*) with the results of Monte-Carlo simulation averaged over 20 random runs at several time points (Fig. 3a). At *t* = 0, simulated and predicted mutant frequencies are constant, since all sites have the same probability of deleterious allele, *f*_in_. Thus, the slope *β* is equal to 0 (blue line). At later times, we observe that the slope increases gradually in time and the frequency of deleterious alleles *f*(*s, t*) depends exponentially on selection coefficient. Apart from some residual fluctuations, our analytical formula (Eq. 1) demonstrates excellent agreement with simulation. Since the sites with possible beneficial mutations with given mutation gain *s* are the sites with deleterious alleles with fitness cost *s*, we confirm that distribution of beneficial fitness effects acquires and maintains an exponential shape in a broad interval of time.

Then, we compared the analytic prediction for the log-slope of DFE *β* (Eq. 2) with simulation, for different values of the average selection coefficient *s*_*av*_ and initial allelic frequency *f*_in_ (Fig. 3b). We observe a good match with the analytic formula that predicts the linear increase of the slope in time at early times. At longer times, *t*>1/*s*_*av*_, our analysis and simulation deviate because the time dependence of the simulated slope becomes slower than linear in time. The results are not very sensitive to initial frequency *f*_in_ or variation of other model parameters (Fig. 3b). Note that, in this regime, although the slope increases more slowly than predicted by Eq. 2, the exponential dependence on mutation cost is conserved. For longer times, the fluctuations increase with time, which is related to strong stochastic effects in the traveling wave regime.

The differences between the prediction of Eq. 2 and simulation at long times are caused by entering the traveling wave regime. In this regime, the wave moves beyond the best-fit sequence present in the initial population due to beneficial mutations (Fig 2f). To predict the slope analytically, we need to account for the effect of beneficial mutations (Good, et al. 2012). Using the analytic result in Eq. 4 derived in *S1 Appendix*, we obtain a good agreement with long-term simulation results (Fig. 3). These results were averaged over several independent simulation runs. Thus, our model of evolution provides a simple explanation for the long-standing puzzle of the exponential DFE (Fig. 1).

### Calculating selection coefficients from a protein or nucleotide sequence set

Our results have an important practical application. They enable us to estimate the relative values of the selection coefficient, *s*, for diverse sites using sequence sets at several points, as long as the system is far from steady-state. Recently, it has been shown that the evolution of influenza in a host population can be mapped on the standard traveling wave theory cited in Introduction with an effective selection coefficient distribution, which depends on immunological and epidemiological properties of the host population (Rouzine and Rozhnova 2018). Hence, the evolution of influenza is amenable to our quasi-stationary or “adiabatic” approximation based on the predictions of traveling wave theory (narrow fitness peak and slow adaptation). When comparing theory with data on influenza below, we work with the average over many subpopulations of influenza collected from different geographic locations around the world.

We start by obtaining a database of aligned sequences of a pathogen or organism obtained at several time points, *t* (at least, two time points). We determine the consensus allele at each aminoacid position, as the most abundant aminoacid variant. Then, we binarize sequences, as follows: we replace each consensus allele with 0, and any minority allele with 1. After binarization, we determine the frequencies of 1 for each and declare it to be the allelic frequency, *f*_*i*_ (*t*). Insertions and deletions are eliminated from analysis. (This technique is appropriate if most diverse sites that have no more than one of two minority variants. Therefore, you cannot use sequences from different species where multiple AA are abundant.)

Based on our analytic results above, the relative value of the selection coefficients at aminoacid position *i* can be estimated from Eq. 6 as

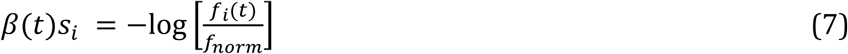

The presence of an additional factor *f*_*norm*_ in Eq. 7 is due to the fact that, in theory, Eqs. 3-6, *f*_*i*_ represents the frequency of less-fit alleles at site *i*, hence, *s*_*i*_ > 0 for all sites by the definition. In real sequences, the best-fit sequence is not known, it is approximated with the consensus sequence. Hence, an anti-consensus allele can be better-fit at some sites, as given by *s*_*i*_ < 0. Since some sites will have negative *s*_*i*_, we need to introduce factor *f*_*norm*_ to account for such sites: they correspond to *f*_*i*_ > *f*_*norm*_.

Factor *f*_*norm*_ is estimated, as follows. Note that the left-hand side in Eq. 7 factorizes into a product of two term: one depends only on time, and another only on site number *i*. This predicts the existence of a fixed point *s*_*i*_<0, independent on time, *t*, which can be used to determine the *norm*alization factor *f*_*norm*_, as follows. For each time, we rank genomic sites in the descending order in *s* and map *i → m*_*i*_ where *i* is the label of an actual site in genome, and *m* is its number after the ranking in *s*. We obtain a monotonous ranked curve, *s*_*rank*_(*m, t*). Then, we find the intersection between curves *s*_*rank*_ (*m, t*) obtained at different times, *t*. Next, we adjust the value of *f*_*norm*_ in Eq. 7 until we obtain *s*_*rank*_ = 0 at the intersection point.

The resulting estimate of β(*t*)*s*_*i*_ from Eq. 7 represents the selection coefficient at site *i* in relative units [1/ *β*]. Further, taking the inverse derivative from each ranked *s* curve, we obtain the distribution density of selection coefficient over non-conserved sites, as given by *g*(*s*), = [*δs*/*δm*]^-1^. Finally, we can re-order the ranked sites back, *i* ← *m*_*i*_ and plot the relative values of selection coefficient, *βs*_*i*_, against their actual aminoacid positions, *i*.

We note in conclusion that frequency *f*_*i*_ (*t*) in Eq. 7, as in Eqs. 2-6, is assumed to be ensemble-averaged, including realizations where site *i* is monomorphic and those where it is polymorphic. Ensemble averaging is approximated in data by combining sequences from different geographic locations.

## Discussion

In summary, we proposed an evolutionary explanation for the exponential DFE of beneficial mutations in terms of the mutation gain in fitness. Using an asexual population model, we predicted a gradual depletion of deleterious alleles with higher fitness costs accompanied by the emergence of a clonal structure after *t* ≈ 1/*s*_*av*_. First, neglecting new mutations, we obtained an exponential dependence of allelic frequency on fitness. The logarithmic slope is equal to time, which corresponds to the virtual absence of linkage effects at early times. The formula is in agreement with Monte-Carlo simulations for early times until *t* ≈ 1/ *s*_*av*_. At longer times, when beneficial mutations become crucial for the generation of new highly fit genomes, we obtained another expression based on the traveling wave theory. Our results confirm the previous work (Pedruzzi, et al. 2018) where an exponential dependence for deleterious allele frequency was predicted using a rather general argument based on the maximum of entropy. This work confirms this result and, moreover, calculates a specific logarithmic slope for the exponential.

Based on the experiments cited in *Introduction*, many models assume an exponential distribution of fitness effects as a starting assumption (Gerrish and Lenski 1998; Good, et al. 2012; Walczak, et al. 2012). Our findings provide an evolutionary justification for this assumption and update these theories by predicting that the distribution is not constant but shrinks in time. However, when mutation selection balance is approached, reverse mutations demolish selection as well as exponential dependence in DFE.

Other groups attempted to explain the universality of the exponential DFE using formal statistical arguments, such as the extreme-value theory (Gillespie 1982; Orr 2003; Joyce, et al. 2008). There are essential differences between this pioneering work and our findings. In the cited work, the aim was to prove an exponential distribution for the raw distribution of selection coefficient among all possible genomic sites, *g(s)*, in the limit of large *s*.

In contrast, we take into account our conclusion that the exponential dependence of DFE on selection coefficient is mostly determined by the evolved occupation numbers of sites *f* (*s, t*) in the broad range of *s* and that *g*(*s*) is a relative slow function of s. Also, the cited approach (Gillespie 1982; Orr 2003; Joyce, et al. 2008) predicts a constant slope of the exponential, while our analysis, simulation, and experimental data prove that it changes in time. Using our result in Eq. 6 backward, we proposed a method to estimate selection coefficients for each diverse site in a sequence sets from two or more time points.

We note that the Fermi distribution for a two-state system with fitness difference, Eq. 6, has emerged in science long before this paper. In 1920s, it was derived by Enrico Fermi for the energy statistical distribution of electrons in equilibrium. For protein configurations and populations in evolutionary equilibrium (steady-state) it has been obtained by (Berg, Willmann, et al. 2004; Sella and Hirsh 2005; Rice, et al. 2015). Both the biological context and the results our work of the present work are quite different from these articles.

In contrast to the cited papers [e.g., Eq. 1 in (Berg, Willmann, et al. 2004)], here we studied a non-equilibrium case. We assumed that the system is very far from equilibrium, *f* ≫ *f*_*eq*_. Eq. 6 does not apply in equilibrium in our evolutionary model, because the presence of new deleterious mutation events, neglected in our derivation for *f* ≫ *f*_*eq*_, mixes the two states and makes the argument invalid. Formula *f*_*eq*_ = *μ*/*s*, must be used instead for large *N*, where *μ* is the mutation rate per site. We have demonstrated this transition to equilibrium for a related model with epistasis (Pedruzzi et al 2018, PLOS CB). Also, the expression for the exponential slope, *β* = −*d*log *f*(*s, t*)/*ds*, differs from the cited papers. Further, it depends on time, which explains the experiments in Fig. 1. This time dependence is absent from the cited papers, because they consider an equilibrium case. The reason why we obtain an quasi-equilibrium result, although with a different “effective temperature” 1/β than in equilibrium, is that the process we studied is adiabatically slow.

To conclude, we demonstrated that the exponential DFE observed in viruses and bacteria is a natural consequence of the process of adaptation. Thus, the new contribution of the present, paper is (i) to explain the existing data in Fig. 1 from the first principles of population genetics, and (ii) to propose a general Method to measure the intrinsic spectrum, *g(s)*, for the adaptation under directional selection. We will consider specific applications to real genomic data elsewhere.

## Materials and Methods

See online Supplement

## Funding

This work has been supported by Agence Nationale de la Recherche grant J16R389 to I.M.R.

## Author contributions

A.B.: Formal analysis, Investigation, Software, Visualization, Writing-original draft

I.M.R.: Conceptualization, Formal analysis, Methodology, Project administration, Supervision, Writing-review & editing

## Competing interests

Authors declare no competing interests.

## Data and materials availability

Code for Fig. 2 and 3 is presented here: https://github.com/irouzine/Barlukova

## Supplementary Materials

Materials and Methods

## Notes

https://github.com/irouzine/Barlukova

### Competing Interest Statement

The authors have declared no competing interest.

## References

Acevedo A, Brodsky L, Andino R. 2014. Mutational and fitness landscapes of an RNA virus revealed through population sequencing. Nature 505:686–690.

Berg J, Lassig M, Wagner A. 2004. Structure and evolution of protein interaction networks: a statistical model for link dynamics and gene duplications. BMC Evol Biol 4:51.

Berg J, Willmann S, Lassig M. 2004. Adaptive evolution of transcription factor binding sites. BMC Evol Biol 4:42.

Brunet E, Rouzine IM, Wilke CO. 2008. The stochastic edge in adaptive evolution. Genetics 179:603–620.

Desai MM, Fisher DS. 2007. Beneficial mutation selection balance and the effect of linkage on positive selection. Genetics 176:1759–1798.

Dutta RN, Rouzine IM, Smith SD, Wilke CO, Novella IS. 2008. Rapid adaptive amplification of preexisting variation in an RNA virus. J Virol 82:4354–4362.

Eyre-Walker A, Keightley PD. 2007. The distribution of fitness effects of new mutations. Nat Rev Genet 8:610–618.

Fisher RA. 1930. The genetical theory of natural selection. Oxford, United Kingdom: Clarendon Press, 1958.

Gerrish PJ, Lenski RE. 1998. The fate of competing beneficial mutations in an asexual population. Genetica 102-103:127–144.

Gillespie JH. 1982. A Randomized Sas Cff Model of Natural-Selection in a Random Environment. Theoretical Population Biology 21:219–237.

Good BH, Desai MM. 2015. The impact of macroscopic epistasis on long-term evolutionary dynamics. Genetics 199:177–190.

Good BH, Rouzine IM, Balick DJ, Hallatschek O, Desai MM. 2012. Distribution of fixed beneficial mutations and the rate of adaptation in asexual populations. Proc Natl Acad Sci U S A 109:4950–4955.

Hallatschek O. 2011. The noisy edge of traveling waves. Proc Natl Acad Sci U S A 108:1783–1787.

Hom N, Gentles L, Bloom JD, Lee KK. 2019. Deep Mutational Scan of the Highly Conserved Influenza A Virus M1 Matrix Protein Reveals Substantial Intrinsic Mutational Tolerance. J Virol 93.

Illingworth CJ, Mustonen V. 2012. Components of selection in the evolution of the influenza virus: linkage effects beat inherent selection. PLoS Pathog 8:e1003091.

Imhof M, Schlotterer C. 2001. Fitness effects of advantageous mutations in evolving Escherichia coli populations. Proc Natl Acad Sci U S A 98:1113–1117.

Joyce P, Rokyta DR, Beisel CJ, Orr HA. 2008. A general extreme value theory model for the adaptation of DNA sequences under strong selection and weak mutation. Genetics 180:1627–1643.

Kassen R, Bataillon T. 2006. Distribution of fitness effects among beneficial mutations before selection in experimental populations of bacteria. Nat Genet 38:484–488.

Keightley PD, Eyre-Walker A. 2007. Joint inference of the distribution of fitness effects of deleterious mutations and population demography based on nucleotide polymorphism frequencies. Genetics 177:2251–2261.

Lee JM, Huddleston J, Doud MB, Hooper KA, Wu NC, Bedford T, Bloom JD. 2018. Deep mutational scanning of hemagglutinin helps predict evolutionary fates of human H3N2 influenza variants. Proc Natl Acad Sci U S A 115:E8276–E8285.

McCandlish DM, Shah P, Plotkin JB. 2016. Epistasis and the Dynamics of Reversion in Molecular Evolution. Genetics 203:1335–1351.

Neher RA, Hallatschek O. 2013. Genealogies of rapidly adapting populations. Proc Natl Acad Sci U S A 110:437–442.

Neher RA, Shraiman BI, Fisher DS. 2010. Rate of adaptation in large sexual populations. Genetics 184:467–481.

Nguyen Ba AN, Cvijovic I, Rojas Echenique JI, Lawrence KR, Rego-Costa A, Liu X, Levy SF, Desai MM. 2019. High-resolution lineage tracking reveals travelling wave of adaptation in laboratory yeast. Nature 575:494–499.

Orr HA. 2003. The distribution of fitness effects among beneficial mutations. Genetics 163:1519–1526.

Pedruzzi G, Barlukova A, Rouzine IM. 2018. Evolutionary footprint of epistasis. PLoS Comput Biol 14:e1006426.

Pedruzzi G, Rouzine IM. 2019a. Epistasis detectably alters correlations between genomic sites in a narrow parameter window. PLoS One 14:e0214036.

Pedruzzi G, Rouzine IM. 2019b. Epistasis detectably alters correlations between genomic sites in a narrow parameter window. PLoS One in press.

Pedruzzi G, Rouzine IM. 2019c. High-fidelity analysis of epistasis predicts primary and secondary drug resistant mutations in influenza., submitted for publication.

Rice DP, Good BH, Desai MM. 2015. The evolutionarily stable distribution of fitness effects. Genetics 200:321–329.

Rouzine IM, Brunet E, Wilke CO. 2008. The traveling-wave approach to asexual evolution: Muller’s ratchet and speed of adaptation. Theoretical Population Biology 73:24–46.

Rouzine IM, Coffin JM. 2005. Evolution of human immunodeficiency virus under selection and weak recombination. Genetics 170:7–18.

Rouzine IM, Coffin JM. 2007. Highly fit ancestors of a partly sexual haploid population. Theoretical Population Biology 71:239–250.

Rouzine IM, Coffin JM. 2010. Multi-site adaptation in the presence of infrequent recombination. Theoretical Population Biology 77:189–204.

Rouzine IM, Coffin JM. 1999. Search for the mechanism of genetic variation in the pro gene of human immunodeficiency virus. J Virol 73:8167–8178.

Rouzine IM, Rodrigo A, Coffin JM. 2001a. Transition between stochastic evolution and deterministic evolution in the presence of selection: general theory and application to virology. Microbiol Mol Biol Rev 65:151–185.

Rouzine IM, Rodrigo A, Coffin JM. (21138142 co-authors). 2001b. Transition between stochastic evolution and deterministic evolution in the presence of selection: general theory and application to virology [review]. Microbiol. Mol. Biol. Rev. 65:151–185.

Rouzine IM, Rozhnova G. 2018. Antigenic evolution of viruses in host populations. PLoS Pathog 14:e1007291.

Rouzine IM, Wakeley J, Coffin JM. 2003. The solitary wave of asexual evolution. Proc Natl Acad Sci U S A 100:587–592.

Sella G, Hirsh AE. 2005. The application of statistical physics to evolutionary biology. Proc Natl Acad Sci U S A 102:9541–9546.

Shah P, McCandlish DM, Plotkin JB. 2015. Contingency and entrenchment in protein evolution under purifying selection. Proc Natl Acad Sci U S A 112:E3226–3235.

Stern A, Bianco S, Yeh MT, Wright C, Butcher K, Tang C, Nielsen R, Andino R. 2014. Costs and benefits of mutational robustness in RNA viruses. Cell Rep 8:1026–1036.

Tsimring LS, Levine H, Kessler D. 1996. RNA virus evolution via a fitness-space model. Phys. Rev. Lett. 76:4440–4443.

Walczak AM, Nicolaisen LE, Plotkin JB, Desai MM. 2012. The structure of genealogies in the presence of purifying selection: a fitness-class coalescent. Genetics 190:753–779.

Wrenbeck EE, Azouz LR, Whitehead TA. 2017. Single-mutation fitness landscapes for an enzyme on multiple substrates reveal specificity is globally encoded. Nat Commun 8:15695.

Yan L, Neher RA, Shraiman BI. 2019. Phylodynamic theory of persistence, extinction and speciation of rapidly adapting pathogens. Elife 8:e44205.

