## Supplement for "The evolutionary origin of the universal distribution of mutation fitness effect"

A. Barlukova , G. Pedruzzi, and I. M. Rouzine*

*Sorbonne Université, Institute de Biologie Paris-Seine, Laboratoire de Biologie Computationnelle et Quantitative, LCQB, F-75004 Paris, France*
*To whom correspondence should be addressed:


**Model.** We consider a haploid population of $N$ binary sequences, where each genome site (nucleotide position) numbered by $i$ =1, 2, …, $L$ carries one of two possible genetic variants (alleles), denoted $K_i$ =0 or $K_i$ =1. The genome is assumed to be very long, $L$ >> 1. Time is discrete and measured in units of population generations. The evolution of the population is simulated using a standard Wright-Fisher model, which includes the factors of random mutation with genomic rate $\mu L$, natural selection, and random genetic drift. Recombination is assumed to be absent. Once per generation, each individual genome is replaced by a random number of its progeny which obeys multinomial distribution. The total population stays constant with the use of the broken-stick algorithm. To include natural selection, the average progeny number (Darwinian fitness) of sequence $K_i$ is set to $e^W$. We consider the simplest case when the fitness effects of mutations, $s_i$ , are additive over sites:

$$W = \sum_{i=1}^{L} s_i\, K_i \tag{8}$$

The reference genome, $\{K_i=0\}$, can be chosen in arbitrary way. For our aim, it is convenient to set it to be the same as the best fit sequence, so that all selection coefficients $s_i$ are negative. Each site $i$ with deleterious allele, $K_i$=1, is a target site for a possible beneficial mutation. Vice versa, a site with the favorable allele, $K_i$=0, can have a deleterious mutation. A more general version of fitness model that accounts for pairwise epistatic interactions is considered in (*2*) and, for macroscopic epistasis, in (*3*). Here we focus on additive contributions of single sites to the fitness landscape.

In our simulations, selection coefficients are chosen randomly at each site from the half-normal distribution

$$g(s) = \frac{2}{s_{av}\pi} \exp\left(-\frac{s^2}{\pi s_{av}{}^2}\right) \tag{9}$$

Here, $s_{av}$ is the average mutation cost in fitness for the initial state.

**Early evolution**. We focus on one genomic site of $L$ sites, whose selection coefficient is assumed to be known and equal to -$s$, where $s$ > 0. The other selection coefficients are assumed to vary following some random distribution with density $g(s)$. We make no assumption regarding the shape of $g(s)$, but assume that distribution of $s$ at different sites is independent. We assume also that initial population has random distribution of alleles among sites with average frequency of less fit alleles denoted by $f_{in}$. Biologically this condition corresponds to a population that has changed suddenly its conditions. First, we will also neglect the effect of new mutations, which simplification is shown to be accurate at early times, when evolution is dominated by decay of standing variation. In the next subsection, we will include the effect of new mutations. Denote by $I_0$ the proportion of all possible sequences having 0 (wild-type) at given site. The proportion of possible sequences having 1 (mutation) at the site denote by $I_1$. Then, the corresponding mutant frequency can be found as ratio

$$f = \frac{I_1}{I_0 + I_1} \tag{10}$$

To obtain frequency in time we account for the action of selection. Selection causes decay of the number of each sequence $\{K_i\}$, by time dependent factor $e^{-t\sum_{i=1}^{L-1} s_i K_i}$. Frequencies $I_0$ and $I_1$ represent the average over all values of $s$ and $K_i$ for all sites

$$I_0 = (1 - f_{in}) \prod_{i=1}^{L-1} \left( \int_0^{+\infty} ds_i \; e^{-t\sum_{i=1}^{L-1} s_i K_i} \sum_{K_i=0}^{1} p(K_i) g(s_i) \right)$$
$$I_1 = f_{in} \; e^{-ts} \prod_{i=1}^{L-1} \left( \int_0^{+\infty} ds_i e^{-t\sum_{i=1}^{L-1} s_i K_i} \sum_{K_i=0}^{1} p(K_i) g(s_i) \right)$$

probabilities of having less-fit and better-fit alleles is $p(1) = f_{in}$ and $p(0) = 1 - f_{in}$, respectively. Then, $I_0$ and $I_1$ can be rewritten by taking exponential term inside of parentheses and separating variables at different sites $i$. The result takes the form

$$f = \frac{e^{-ts} f_{in}}{1 - f_{in} + e^{-ts} f_{in}} \tag{11}$$

**Traveling wave regime.** In the traveling wave regime, which starts around $t > 1/s_{av}$, beneficial mutations have to be included into consideration because they create new best-fit genomes of the traveling wave (*4-7*). Let $t_0$ be the characteristic time of the beginning of traveling wave regime and $\pi(s)$ be the fixation probability of beneficial mutations. In this regime, most of deleterious alleles are located at uniformly deleterious sites (Fig. 2, yellow columns). Hence, their loss occurs mostly due to fixation of new beneficial alleles at these sites. Then, the dynamic equation for the frequency of deleterious alleles for $t > t_0$ , $t_0 \sim 1/s_{av}$ , can be written as follows

$$\frac{\partial f(s,t)}{\partial t} = -\mu N \pi(s) f(s,t) \tag{12}$$

The initial condition for Eq. 8 can be obtained from the estimate of $f$ in the initial time interval where selection of pre-existing genomes is the dominant process. From Eq. 7

$$f(s, t_0) \approx f_{in} e^{-t_0 s}$$

The solution of Eq. 8 with these initial conditions is given by Eq. 3 in the Main text. Thus, the problem is reduced to the result of the previous work (*8*) for the fixation probability of mutations $\pi(s)$

$$\pi(s) \sim e^{-\frac{s^2}{2v}} \left( \frac{e^{-s\frac{x_C}{v}} - 1}{s} \right) + \frac{e^{\frac{x_C^{\,2}}{2v}}}{v x_C} \int_{x_C}^{\infty} dx \, x e^{-\frac{(x-s)^2}{2v}}, \quad \pi(0) \approx \frac{1}{N} \tag{13}$$

Here, $v$ is the rate of adaptation, $x_c$ is the characteristic value of fitness. Parameters $v$ and $x_c$ can be found numerically from two transcendental equations

$$2 = \frac{U_b}{\sigma} \sqrt{\frac{2\pi\sigma^2}{v}} \left[ 1 + \frac{\sigma}{x_C} + \frac{v}{\sigma x_C - v} \right] e^{\frac{(x_C - \frac{v}{\sigma})^2}{2v}} \tag{14}$$

$$1 = N U_b \left[ \frac{x_C^2}{v} - 1 + \frac{2 x_C \sigma}{v} + \frac{2\sigma^2}{v} \right] e^{-\frac{1}{\sigma}\left(x_C - \frac{v}{2\sigma}\right)} \tag{15}$$

where $U_b$ is the probability of beneficial mutation per genome per generation. Thus, the time evolution of the frequency of deleterious alleles can be found numerically using Eqs. 3, 9, 10, 11. We note that parameter $t_0$ in Eq. 3 is unknown, since Eq. 3 is the asymptotic expression at large times, $t \gg 1/s_{av}$ To connect two regimes of evolution we find optimum value of $t_0$, such that simulated and analytical curves for the slopes are at the best match. The value of $\sigma$ is the average effect of beneficial mutation for the sites that can have such a mutation and, hence, is equal to $1/\beta$, where $\beta$ found from self-consistent condition given by Eqs. 2 and 3. Note that Eq. 3 predicts an exponential decay at small to moderate $s$; at large $s \sim 1$, it predicts a faster decay with $s$.


1. G. Pedruzzi, I. M. Rouzine, High-fidelity analysis of epistasis predicts primary and secondary drug resistant mutations in influenza., *submitted for publication*, (2019).
2. G. Pedruzzi, A. Barlukova, I. M. Rouzine, Evolutionary footprint of epistasis. *PLoS Comput Biol* **14**, e1006426 (2018).
3. B. H. Good, M. M. Desai, The impact of macroscopic epistasis on long-term evolutionary dynamics. *Genetics* **199**, 177-190 (2015).
4. I. M. Rouzine, J. Wakeley, J. M. Coffin, The solitary wave of asexual evolution. *Proc Natl Acad Sci U S A* **100**, 587-592 (2003).
5. M. M. Desai, D. S. Fisher, Beneficial mutation selection balance and the effect of linkage on positive selection. *Genetics* **176**, 1759-1798 (2007).

6. I. M. Rouzine, J. M. Coffin, Highly fit ancestors of a partly sexual haploid population. *Theor Popul Biol* **71**, 239-250 (2007).
7. R. A. Neher, B. I. Shraiman, D. S. Fisher, Rate of adaptation in large sexual populations. *Genetics* **184**, 467-481 (2010).
8. B. H. Good, I. M. Rouzine, D. J. Balick, O. Hallatschek, M. M. Desai, Distribution of fixed beneficial mutations and the rate of adaptation in asexual populations. *Proc Natl Acad Sci U S A* **109**, 4950-4955 (2012).
